# Translational approach to social isolation during a global pandemic: Hippocampal somatic mutation

**DOI:** 10.1101/2023.06.06.542200

**Authors:** Bomee Lee, Yuri Seo, Sohee Jung, Soojung Im, Hyung Jun Choi, Jae Nam Bae, Yangsik Kim

**Author notes:** Corresponding author: Yangsik Kim, MD, PhD. Department of Psychiatry, Inha University Hospital, Incheon, South Korea.

## Abstract

The coronavirus disease 2019 (COVID-19) pandemic has attributed to stress not only by the infection itself but also by social isolation owing to self-quarantine and social distancing. Stress has adverse effects on the mental health and chronic medical diseases; the potential of stress-induced somatic mutations in the brain to cause psychiatric disorders is being studied. Here we conducted behavioral studies, protein expression studies, single-nucleus sequencing (snRNAseq), and whole-genome sequencing (WGS) of the hippocampus of mice that underwent early maternal separation and social isolation, and a demographic study of community populations who had been self-quarantined owing to COVID-19 exposure to investigate the link between somatic mutations and stress due to social isolation. The demographic study demonstrated more negative mental health findings among individuals who live alone or are single. Mice subjected to early maternal separation and social isolation demonstrated increased anxiety-like behaviors and stress-related corticotropin-releasing hormone receptor 1, and neurogenesis-related sex-determining region Y-box 2 and doublecortin expression. In snRNA-seq, differences, such as transthyretin increase, were observed in the maternal separation group, and somatic mutations, including insertion in the intron site of Tmem267, were observed in the social isolation group on WGS. The results of this study suggest that stress, such as social isolation, can cause changes at the genetic level, as well as behavioral and brain protein changes.

## INTRODUCTION

The unprecedented COVID-19 pandemic has forced the world to close doors and maintain social and personal isolation. In particular, isolation by self-quarantine and physical distancing has adversely affected mental health conditions, such as depression and anxiety, which are expected to continue to have an impact even now despite the waning impact of COVID-19.(1–3) According to the World Health Organization, depression and anxiety increased by 25% worldwide in the first year of the COVID-19 outbreak, with social isolation, loneliness, sadness, and economic problems contributing to unprecedented stress, mental health challenges, and exposure to existing health inequalities.(4–6) Mental health challenges attributed to COVID-19 are particularly pronounced among women,(6) adolescents,(7) and populations with medical comorbidities.(7) Such mental health problems appeared not only during the COVID-19 pandemic, but also during the Spanish flu and H5N1 pandemics.(8, 9) Therefore, =ensuring that individuals are well prepared for the mental health challenges associated with the pandemic is crucial.

Social isolation is a well-known cause of stress,(10) which affects not only humans but also animals forming social groups, such as mice.(1, 11) Social isolation is associated with depression, anxiety, and several aspect of mental health, especially social isolation in early life is known to affect neurotransmitters, such as the corticotropin releasing hormone (CRH), dopamine, norepinephrine, serotonin, glutamate, and gamma-aminobutyric acid (GABA).(1)

Stress has a strong relationship not only with mental health but also with medical diseases.(12) It is also associated with chronic diseases, such as diabetes, hypertension, and cancer. In particular, since stress and toxic substances present in the environment can affect somatic mutation, somatic mutation research has recently attracted attention in cancer research.(13–15) Currently, studying somatic mutations in the brains of individuals with psychiatric conditions other than Alzheimer’s disease, in which HPA axis-related factors are induced by stress, is warranted.(16)

In this study, the behavioral changes in humans and mice attributed to social isolation were evaluated, and changes in the hippocampal protein expression in mice were investigated. Moreover, we performed single-cell RNA and whole-genome sequencing of the hippocampus to investigate the genetic changes caused by social isolation.

## RESULTS

### Demographic factors of self-quarantined individuals owing to COVID-19

Statistical analysis was conducted based on the demographic factors of the uninfected population who were self-quarantined owing to COVID-19 (Table 1). We compared the population according to sex, marital status, and residential status. There were differences in depression (female [F] 3.2 vs. Male [M] 4.2, *p* = 0.002), anxiety (F 2.0 vs. M 2.6, *p =* 0.02), somatic symptoms (F 5.3 vs. M 8.0, *p* <0.001), suicide (F 2.5 vs. M 2.2, *p =* 0.004), sleep (F 6.2 vs. M 7.6, *p =* 0.003), and stress perception (F 14.8 vs. M 15.9, *p =* 0.009), and PTSD symptoms (F 8.3 vs. M 10.5, *p* = 0.04).

**Table 1.** Self-reported findings on mental health of individuals self-quarantined owing to coronavirus disease 2019 and differences according to variables including Gender, marriage, and family.

|  | Mean<br>or n | SD or % |
| --- | --- | --- |
| **Demographics** |  |  |
| Sex, female | 311 | 55.6% |
| Age | 39.4 | 12.93 |
| Education | 14.2 | 2.78 |
| Marriage status, alone | 246 | 44.1% |
| Family status, alone | 146 | 26.2% |
| **Self-Report Scale Score** |  |  |
| Depression | 3.8 | 3.95 |
| Anxiety | 2.3 | 3.32 |
| Somatic symptoms | 6.8 | 4.94 |
| Suicide | 2.3 | 1.41 |
| Sleep | 7.0 | 5.86 |
| Stress | 15.4 | 5.07 |
| PTSD | 9.6 | 12.72 |
| Self-esteem | 31.5 | 5.27 |

| **Sex** | female 311 |  | male 247 |  |  |  |
| --- | --- | --- | --- | --- | --- | --- |
|  | Mean | SD | Mean | SD | p.value | t |
| Age | 39.9 | 13.10 | 38.9 | 12.80 | 0.36 | 0.92 |
| Education | 14.1 | 3.01 | 14.3 | 2.58 | 0.31 | -1.02 |
| Depression | 3.2 | 3.73 | 4.2 | 4.06 | 0.002 | -3.19 |
| Anxiety | 2.0 | 3.03 | 2.6 | 3.51 | 0.02 | -2.31 |
| Somatic symptoms | 5.3 | 4.39 | 8.0 | 5.03 | <0.001 | -6.64 |
| Suicide | 2.5 | 1.41 | 2.2 | 1.39 | 0.004 | 2.89 |
| Sleep | 6.2 | 5.33 | 7.6 | 6.18 | 0.003 | -2.95 |
| Stress | 14.8 | 4.57 | 15.9 | 5.39 | 0.009 | -2.63 |
| PTSD | 8.3 | 11.50 | 10.5 | 13.55 | 0.04 | -2.05 |
| Self-esteem | 31.6 | 5.48 | 31.4 | 5.10 | 0.60 | 0.52 |

| **Marriage** | Single 246 |  | Married 312 |  |  |  |
| --- | --- | --- | --- | --- | --- | --- |
|  | Mean | SD | Mean | SD | p.value | t |
| Age | 32.2 | 12.12 | 45.0 | 10.54 | <0.001 | -13.52 |
| Education | 13.8 | 3.04 | 14.5 | 2.52 | 0.008 | -3.77 |
| Depression | 4.2 | 4.16 | 3.4 | 3.74 | 0.02 | 2.15 |
| Anxiety | 2.4 | 3.52 | 2.2 | 3.15 | 0.49 | 0.45 |
| Somatic symptoms | 6.6 | 5.09 | 7.0 | 4.81 | 0.44 | -1.01 |
| Suicide | 2.1 | 1.36 | 2.5 | 1.43 | 0.001 | -3.18 |
| Sleep | 7.3 | 6.25 | 6.7 | 5.53 | 0.19 | 0.95 |
| Stress | 15.6 | 5.53 | 15.2 | 4.68 | 0.45 | 0.53 |
| PTSD | 9.4 | 13.07 | 9.7 | 12.45 | 0.83 | -0.34 |
| Self-esteem | 30.8 | 5.88 | 32.0 | 4.67 | 0.006 | -2.49 |

| **Residence** | Alone 146 |  | Together 412 |  |  |  |
| --- | --- | --- | --- | --- | --- | --- |
|  | Mean | SD | Mean | SD | p.value | t |
| Age | 35.6 | 11.16 | 40.7 | 13.26 | <0.001 | -4.47 |
| Education | 14.6 | 2.38 | 14.1 | 2.90 | 0.03 | 2.20 |
| Depression | 4.2 | 4.31 | 3.6 | 3.81 | 0.17 | 1.39 |
| Anxiety | 2.7 | 3.79 | 2.2 | 3.13 | 0.10 | 1.67 |
| Somatic symptoms | 6.8 | 5.19 | 6.8 | 4.85 | 0.91 | -0.11 |
| Suicide | 2.2 | 1.38 | 2.4 | 1.42 | 0.09 | -1.69 |
| Sleep | 8.1 | 6.42 | 6.6 | 5.60 | 0.009 | 2.64 |
| Stress | 15.7 | 5.28 | 15.3 | 5.00 | 0.45 | 0.76 |
| PTSD | 10.2 | 12.95 | 9.3 | 12.64 | 0.47 | 0.72 |
| Self-esteem | 30.6 | 5.64 | 31.8 | 5.11 | 0.03 | -2.16 |

Abbreviation: PTSD, post-traumatic stress disorder; SD, standard deviation.

Marital status was divided into single (single, separated, and divorced) and married (married and cohabitating). There were differences in the age (single [S] 32.2 vs. married [M] 45.0, *p* < 0.001), year of education (S 13.8 vs. M 14.5, *p* = 0.008), depression (S 4.2 vs. M 3.4, *p* = 0.02), suicide (S 2.1 vs. M 2.5, *p* = 0.001), and self-esteem (S 30.8 vs. M 32.0, *p* = 0.006).

According to the residence status, the status of living alone (A) and living together (T) (including both family and non-family) were compared. There were differences in the age (A 35.6 vs. T 40.7, *p <* 0.001), year of education (A 14.6 vs. T 14.1, *p =* 0.03), sleep (A 8.1 vs. T 6.6, *p =* 0.009), and self-esteem (A 30.6 vs. T 31.8, *p =* 0.03). There were no statistical differences between variables such as sex and marriage, sex and residence, and marriage and residence.

Overall, male, being single, and living alone tended to have negative mental health outcomes in self-quarantined people; we further investigated the effects of social isolation on the brain and mental health through animal and genomic studies.

### Behavioral and protein expression measurements to investigate the effects of early maternal separation and social isolation in mice

Based on the demographic results, we conducted an isolation test on mice to evaluate the biological mechanism of the effect of social isolation; early maternal separation was used to hypothesize the factors for individual vulnerability along with social isolation in adult mice (Figure 1A).

**Figure 1.**
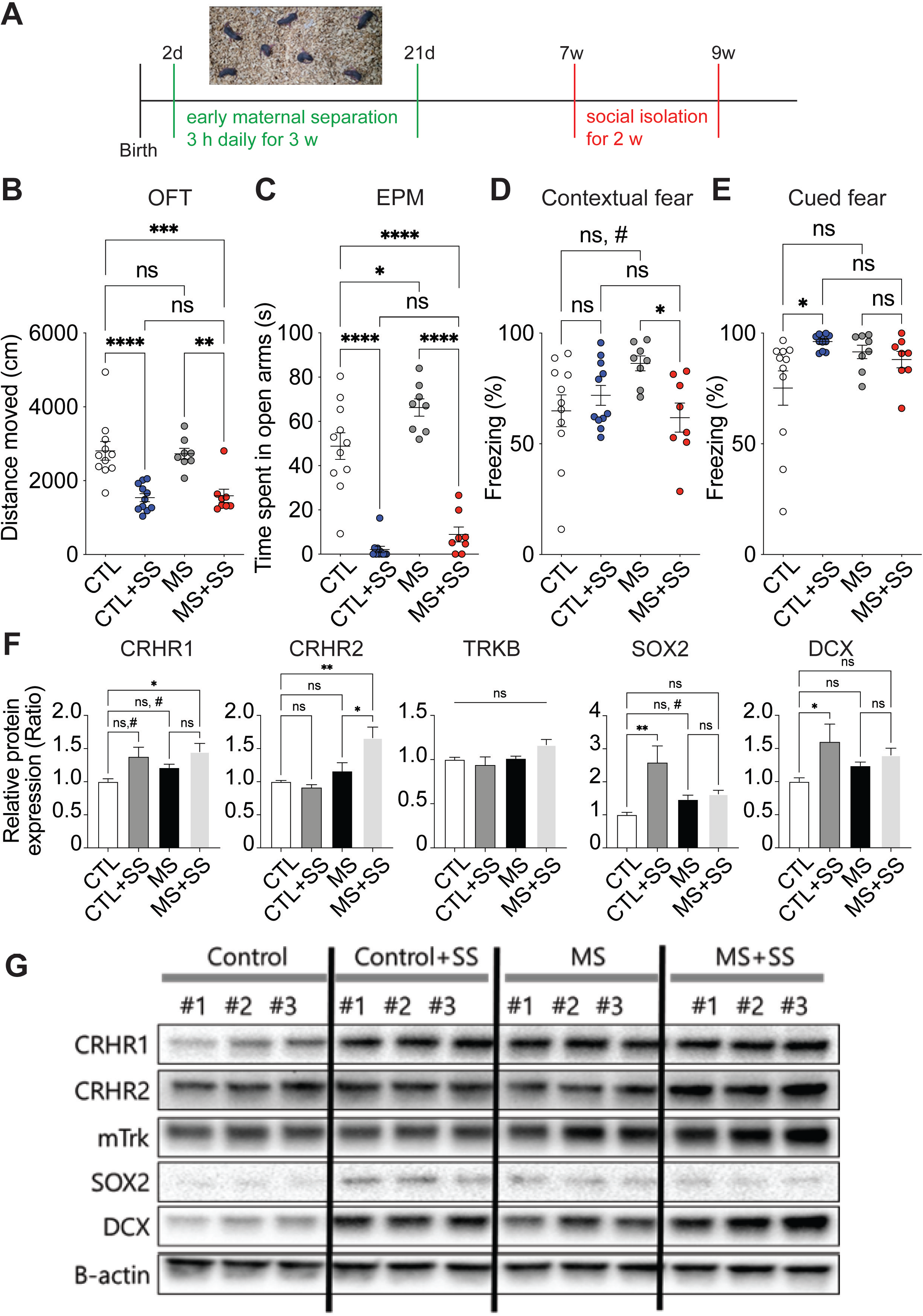
Changes in the behavior and protein expression in maternal separation and socially isolated mice. (**A**) Description of the mouse behavior experiment method. (**B–E**) There was no difference in the locomotion (OFT) between the control group (CTL) and the maternal separation group (MS), and the two groups that underwent social isolation (social isolation after control [CTL+SS] and social isolation after maternal separation [MS+SS]) decreased when compared to CTL and MS, respectively. Anxiety-like behavior (EPM) showed that the time spent in the open arm increased in the MS compared to CTL, and the time spent in the open arm significantly decreased in the CTL+SS and MS+SS compared to the CTL and MS. Considering fear-related behavior, the MS+SS showed a decrease in freezing in the contextual fear test compared to the MS, and an increase in freezing in the cued fear test in the CTL+SS compared to the CTL. The MS showed increased freezing in the contextual fear test compared to the CTL in the t-test (n = 11 mice [CTL], 8 [CTL+SS], 11 [MS], 8 [MS+SS]). (**F**) Compared to the CTL, the MS+SS showed significant increase in the CRHR1 and CRHR2, which are related to stress and hypothalamic-pituitary-adrenal axis. Moreover, neurogenesis-related SOX2 and DCX demonstrated an increase in the social isolation group (CTL+SS). TRKB did not show a difference between the groups. In the t-test, CRHR1 increased in the MS and CTL+SS, and DCX increased in the CTL+SS group. It was standardized based on the protein expression ratio of the hippocampi of the control mice group (n = 9 mice [CTL], 9 [CTL+SS], 9 [MS], 9 [MS+SS], One-way analysis of variance, **** p<0.0001, *** p<0.001, ** p<0.01, * p<0.05; Student’s t-test, # p<0.05, *post hoc* Tukey). (**G**) Representative images of immunoblots in the mouse hippocampus. CRHR1, Corticotropin-releasing hormone receptor 1; CRHR2, Corticotropin-releasing hormone receptor 2; TRKB, Tropomyosin receptor kinase B (mouse); SOX2, SRY (sex-determining region Y)-box 2; DCX, Doublecortin.

Locomotion observed through the open field test did not demonstrate a difference between the control and maternal separation groups, whereas locomotion measured after social isolation demonstrated a difference within the group but not show a difference between the groups (Figure 1B). In the elevated plus maze test to measure anxiety behavior (Figure 1C), the time spent in the open arms in the maternal separation group increased compared to the control group, and after social isolation, a difference was observed within the group, but not between the groups. In the fear conditioning test using foot shock (Figure 1D and E), the maternal separation group demonstrated increased freezing compared to the control group, and the sound cue-induced freezing behavior was different from that of the control group (control vs. control following social isolation) but did not show a difference within the maternal separation group.

We observed the effects of maternal separation and social isolation on the expression of proteins in the hippocampi of the mice (Figure1 F and G). Immunoblotting was performed, focusing on the HPA axis and neurogenesis abnormalities caused by stress. Corticotropin-releasing hormone receptor 1 (CRHR1) expression was increased in the maternal separation group compared to that in the control group, and maternal separation and social isolation demonstrated an increased pattern compared to that in the control group, even upon multiple comparisons. CRHR2 expression was higher in the maternal separation and social isolation group than in the control group. There was no difference between the groups in the tropomyosin receptor kinase B (TrkB) that Brain Derived Neurotrophic Factor binds to; SOX2, a neural stem cell marker, and doublecortin (DCX), a neuro progenitor cell marker, were increased in the MS group compared to the control group. Except for MAP2, there were no differences in the major downstream signaling pathways and associated proteins of CRHR1, such as GRIN2B, mTOR, p-mTOR, ERK, and p-ERK (Supplementary Figure 1).

### Single-nucleus RNA sequencing (snRNAseq) was performed to analyze single-cell transcriptome changes in the hippocampus of mice subjected to maternal separation and social isolation

We performed genomic analysis to identify single-cell-level transcriptome changes in the mouse brain, specifically in the hippocampus, which is directly affected by stressors, such as social isolation and maternal separation. SnRNAseq analysis was performed to observe somatic mutations and transcriptional changes resulting from the effect of stress in each cell.

snRNAseq and analysis were performed on the control group (CTL), maternal separation group (MS), and social isolation after maternal separation group (SS) (three mice in each group). Each group showed a similar cell distribution (Supplementary Figure 2A and B). In the CTL; neurons accounted for 92.1%, astrocytes for 4.9%, and oligodendrocytes for 2.3% (Figure 2B and Supplementary Figure 2C). Each cluster was classified according to the transcript expression (Supplementary Figure 2D and E).

**Figure 2.**
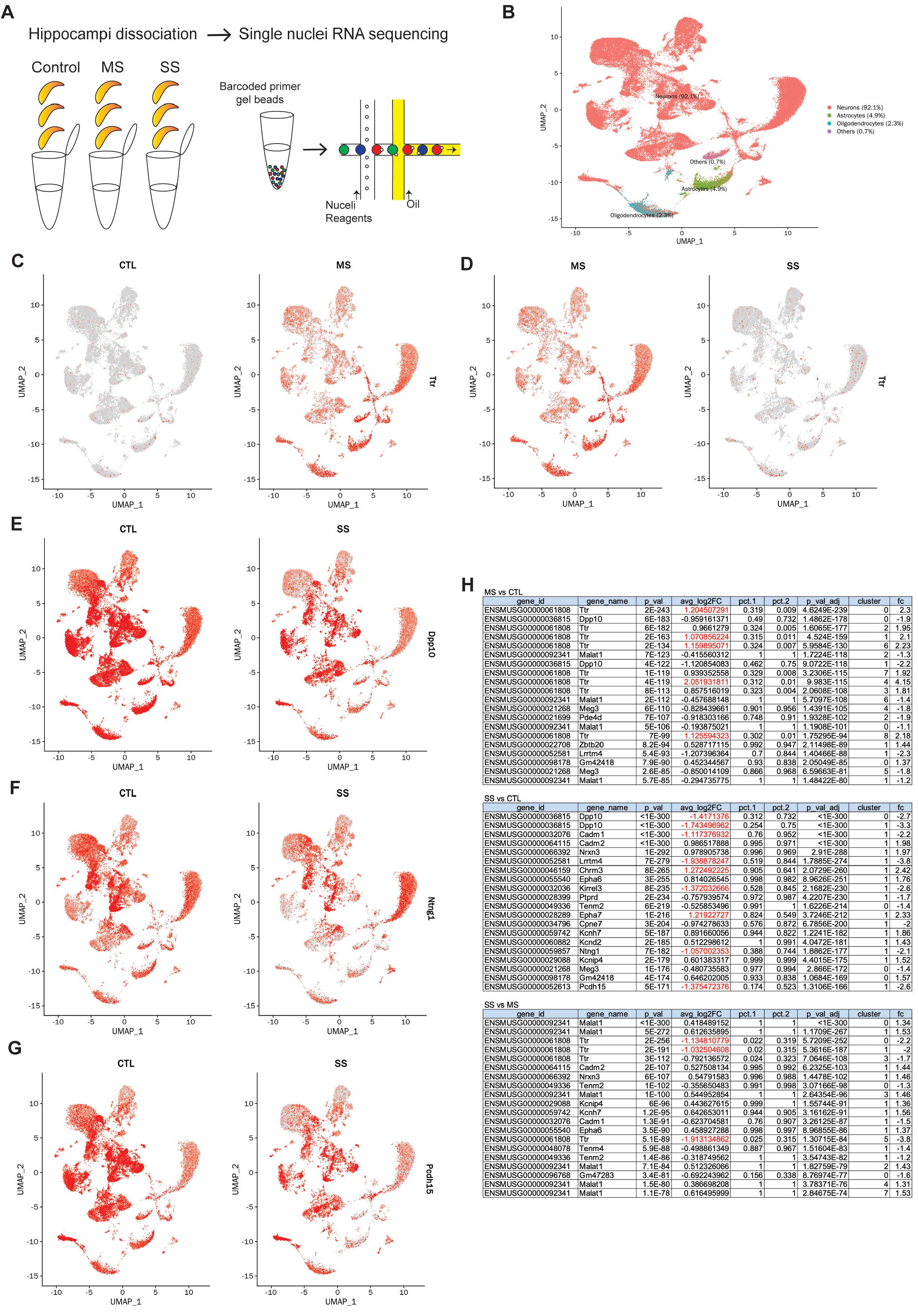
Single nucleus transcriptome changes in mice with maternal separation and social isolation. (**A**) Schematic illustration of single nucleus RNA sequencing. (**B)** Identified cell types by Seurat analysis from single nucleus RNA sequencing in mouse hippocampi and cell type distribution between the groups. **(C–F**) FeaturePlot (above) and VlnPlot (below) showing differentially expressed genes between groups at the cell level for each cluster. In the maternal separation group (MS), the expression level of transthyretin (*Ttr)* increased more than twice as much as that of the control group (CTL) (**B**). In the maternal separation with social isolation group (SS), the *Ttr* decreased more than twice as compared to the MS (**C**). In the SS, compared to the CTL, *Dipeptidyl peptidase 10 (Dpp10), Netrin G1 (Ntng1),* and *Protocadherin 15 (Pchd15)* decreased more than twice as much (**D–F**). (**G**) Lists showing the top 20 genes by group (n = The hippocampi of 3 mice were included in 1 assay each in the CTL, MS, and SS; adjusted p-values were corrected by the Bonferroni correction.).

Differences in Differentially Expressed Genes (DEGs) analyses were compared by group, and the expression of transthyretin (*Ttr*)) increased in the MS compared to that in the CTL; increased expression was observed in all clusters, including neurons, astrocytes, and oligodendrocytes (Figure 2C and H). Between MS and SS, an increase in *Ttr* was observed in the MS (Figure 2D). In the SS compared to the CTL, decreases in *Dipeptidyl peptidase 10 (Dpp10), Netrin G1 (Ntng1),* and *Protocadherin 15 (Pchd15)* were observed (Figure 2E-G).

### Somatic mutations revealed by WGS in the hippocampus of mice with maternal separation and social isolation

We observed stress-induced DNA-level somatic mutations in the hippocampi of mice subjected to stressors, such as maternal separation and social isolation. WGS was performed in the heart of each mouse at approximately 30×depth and hippocampus at approximately 100×depth. In addition, the WGS results for the heart and hippocampus were compared, and the results for each individual were compared according to the group.

First, in the WGS results of the heart and hippocampus, a difference in the total variants, including insertions, deletions, and single nucleotide polymorphisms (SNPs) was observed; however, no difference was observed in each group (Figure 3A and B). This could be attributed to the differences in the organs; however, the effect of sequencing sensitivity according to sequencing depth coverage should also be considered.

**Figure 3.**
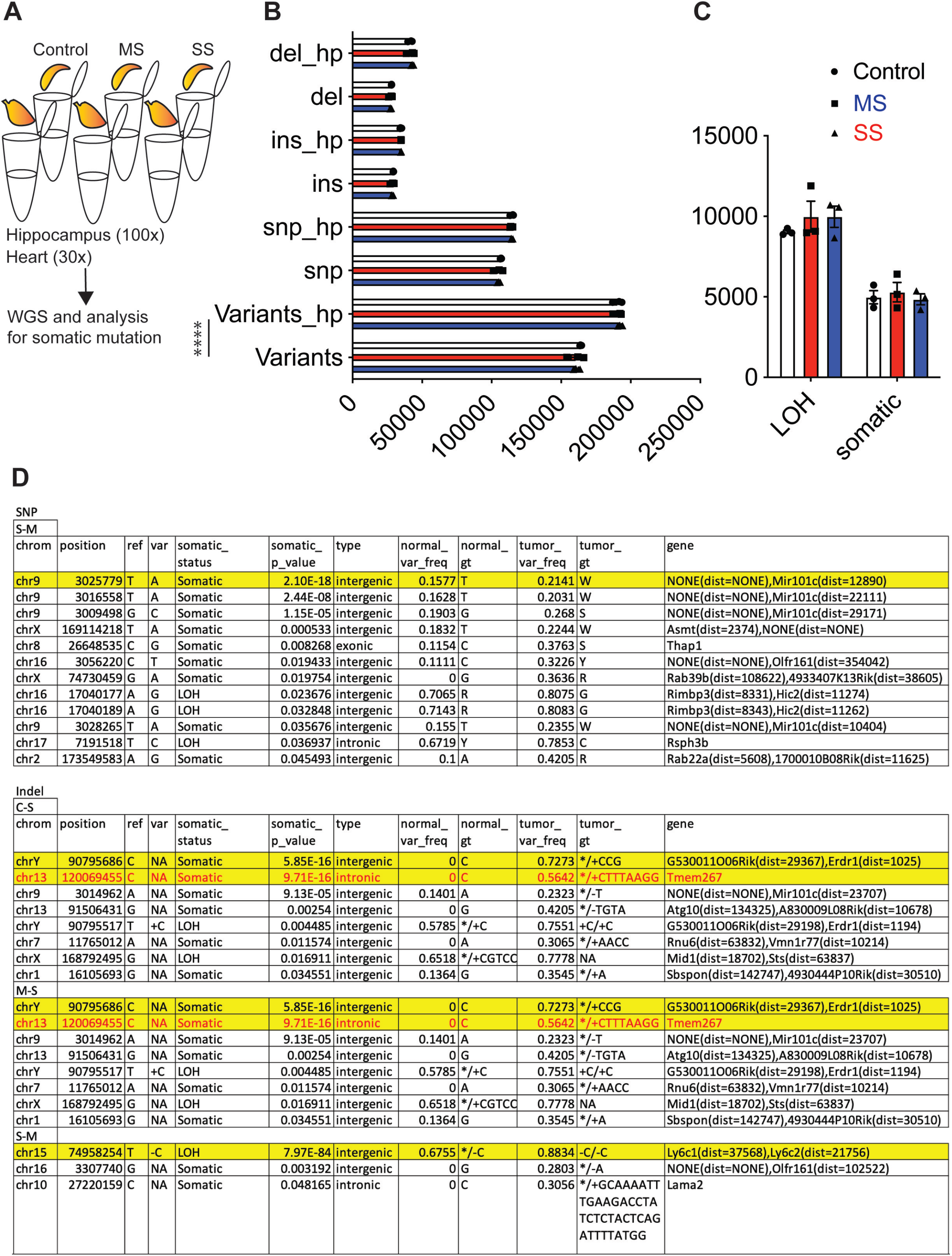
Results of whole-genome sequencing of mice with maternal separation and social isolation based on the differences in the genetic changes in the hippocampus and heart. (**A**) Schematic illustration of whole-genome sequencing to investigate somatic mutations of the hippocampus compared to the heart. (**B**) Total variants including single nucleotide polymorphism (SNP), insertion (ins), and deletion (del) were increased greatly in the hippocampus (100× depth coverage) than in the heart (30× depth coverage) of mice. (**B**) No differences were observed between the groups in loss of heterogeneity (LOH) and somatic mutation. (**C**) Lists representing somatic mutations including SNPs and Indels among groups (n = The hippocampi of 3 mice each in the CTL, MS, and SS)

Although the genome-wide significance *p*-value threshold of 5 × 10^−8^ has become a standard for common-variant genome-wide association studies for lower frequency variants, we interpreted our results for 1 × 10^−9^ or higher as significant for more stringent statistics. Among the SNPs (Figure 3C), a significant difference was observed in the chromosome 9 3025779 intergenic position (T -> A) between the MS and SS. In insertion and deletion (Indel), compared to the CTL, the intergenic somatic mutation at the chr Y 90795686 position (CCG insertion) and the somatic mutation in the intronic region of the *Tmem267* gene at the chr 13 120069455 position (CTTTAAG insertion) demonstrated significant differences in SS. Between the MS and SS, the above chr Y 90795686 and *Tmem267* intronic somatic mutations appeared in the SS compared to the MS, and loss of heterogeneity of the intergenic region at chr 15 74958254 (C deletion) was observed in the MS compared to SS. Somatic mutation profiles for each mouse are shown in Supplementary Table 1.

Using WGS, somatic mutations in the MS and SS were confirmed and compared with those in the CTL. In particular, *Tmem267* revealed somatic mutations in the intron region that could directly affect the transcriptome. Intergenic somatic mutations were also observed.

## DISCUSSION

In this study, socially isolated groups of individuals who self-quarantined owing to COVID-19 exposure demonstrated worse results in terms of depression, anxiety, and mental health measurements. In addition, anxiety-related behaviors increased in mice subjected to early maternal separation and social isolation. The protein expression of CRHR1, a stress-related CRH receptor, and neurogenesis-related SOX2 and DCX increased in the hippocampi of socially isolated mice. Single-nucleus RNA-sequencing revealed that mice subjected to early maternal separation and social isolation showed transcriptome changes, including increased *Ttr1* expression and decreased *Dpp10*, *Ntng1*, and *Pchd15* expression. Finally, multiple somatic mutations, including the *Tmem267* intronic mutation in the MS+SS, were observed in the hippocampi of the mice.

Social isolation, one of the major stressors in this study, is related to the deterioration of human mental health, such as depression and anxiety. Animal experiments confirmed changes in anxiety behavior and stress-related proteins in the hippocampi following social isolation. In addition, somatic mutations through WGS and transcriptome changes upon single-nucleus RNA sequencing could be induced by isolation stress.

In a study conducted on somatic mutations in the human brain related to psychiatric disorders,(17–19) 131 postmortem brains of patients with schizophrenia, Tourette’s syndrome, autism spectrum disorder, and a healthy control group were analyzed using WGS at 200x coverage.(17) A total of 20–60 SNV were observed, and more than 101 hypermutable brains were distributed according to age rather than disease classification. In a study using whole exome sequencing (WES) of 500× or more coverage of postmortem brains in the Alzheimer’s disease and healthy control groups, no difference was observed between the groups; however, an increase in the mutations with age was observed, and 22 somatic mutations per year were predicted in the hippocampal formation(20) (40 somatic mutations per year were predicted in the dentate gyrus in the healthy control group only in the study).(21) In a study of patients with schizophrenia known to have a heritability of approximately 80%,(18) eight genetic mutations of a total of 106 somatic deletions were reported in three patients compared to two healthy controls.(22) Another study in patients with schizophrenia demonstrated an increase in long interspersed nuclear element-1, which is known to be increased in Rett syndrome.(23) In addition, in a recent WES study, no difference was observed in the number of somatic mutations between the schizophrenia patient group and control group; however, mutations in the SCZ-associated pathways, including *GRIN2B*, were reported to be more prevalent in the schizophrenia group.(24) In this study, similar to previous studies, no difference was observed in the number of somatic mutations in the hippocampi of the MS, MS+SS, and CTL mice. In particular, the *Tmem267* intron mutation was commonly observed in the MS + SS of WGS.

In a study of stress-related somatic mutations,(25–27) an RNA-seq study using the blood of soldiers with post-traumatic stress disorder demonstrated 187 and 442 cases of high and moderate severity of transcriptional abnormalities, respectively, including inflammation-related genes.(28) Prenatal stress results in oxidative damage of the mitochondrial DNA in the hippocampus of newborn mice(29). Post-mortem brain studies of 63 psychiatric conditions, including schizophrenia, major depressive disorder, and bipolar affective disorder, have also demonstrated mitochondrial DNA damage.(30)

As described above, inflammation and oxidative stress affect the mechanisms underlying stress-related somatic mutations.(31–33) Studies have been conducted on the possibility of genetic damage attributed to inflammatory and oxidative damage caused by environmental factors, including stress in neurons, which are the main cells of the central nervous system having a very low replication rate.(29, 31, 33) Several studies have reported genetic variants in mental disorders, such as schizophrenia, major depressive disorder, and bipolar affective disorder, that first appear in early adulthood. However, the results are inconsistent, and genetic findings are spread across multiple diagnoses rather than a single diagnosis.(18, 34, 35) This could be related to the underlying mechanisms by which somatic mutations occur randomly at various developmental stages, such as increased mutations in long genes and active transcription.(32, 34, 36, 37) In the future, research with more emphasis on the genetic changes caused by environmental factors, especially stress, is warranted.

Individual variations in stress in mice were identified upon analyzing of the results of all the experiments. Therefore, to observe a clearer difference, molecular and genetic analyses should be performed after selecting mice more susceptible to stress following behavioral experiments in future studies. In addition, this study only covered 100-fold coverage in WGS. Ideally, a coverage of 200-fold or higher is considered more sensitive to somatic mutations; thus, higher coverage is warranted. In addition, since only demographic data existed in the human data in this study, the results of this study should be reconfirmed if blood and hippocampus samples of the participants could be obtained and analyzed from a well-established biobank with clinical information, such as stress measurements.

The human study was based on demographic data, including depression, anxiety, and other mental health issues, among individuals self-quarantined owing to COVID-19 exposure. The study findings are valuable since the study yielded social experimental data obtained in a situation where individuals were artificially isolated because of COVID-19. In addition, early maternal separation and social isolation were objectively replicated through animal experiments, and changes in stress were observed at various levels, such as behavioral experiments, protein expression, snRNAseq, and WGS. Somatic mutations resulting from maternal separation and social isolation have also been identified.

In this study, changes in mental health, such as depression and anxiety, in the general public owing to social isolation during the COVID-19 pandemic and the underlying mechanism were explored through animal experiments, snRNAseq of the hippocampus, and 100×WGS. This study provides evidence that genetic-level changes, such as somatic mutations, could be attributed to stress such as social isolation.

## METHODS

### Animals and Experimental Design

C57BL/6J male mice were purchased from DBL Co. (South Korea) and provided with free access to food and water and four to five animals were housed per cage under a 12-hour light-dark cycle. We included only male mice in this study because demographic studies showed more negative mental health in males. All the mice were bred and maintained in accordance with the requirements of the Animal Research at National Center for Mental Health (NCMH), and all procedures and methods were approved by the Committee of Animal Research at NCMH (IACUC No: NCMH-2005-001-003-01).

C57BL/6J pregnant mice were supplied on the 12th day of TP (Time Pregnant), acclimatized for about a week, and then used in the experiment. The maternal separation group was separated from the mother and other cages for 3 hours each for 3 weeks from the 2nd day after birth, and the adult separation group was separated from the mother and social isolation group at 8 weeks of age by individual cages for 2 weeks. Mice were randomly assigned to either the maternal separation or control groups, and after reaching adulthood, they were randomly assigned to the social isolation group.

### Open-field test

Mice were placed in the center region of an open-field box (45 x 45 x 45 cm), and the locomotor activity in the open field arena was measured for the first 5 minutes. Behavioral tests were recorded and analyzed using SMART 3.0^®^ tracking software (Panlab Harvard Apparatus, Barcelona, Spain).

### Elevated plus-maze test

An elevated-plus maze made of gray acrylic with four arms, each 25-cm long and 2.5-cm wide, positioned 16-cm above the ground, was used for measuring anxiety-like behavior. Light intensity in the closed arms was approximately 0 lux. The test was initiated by placing the mouse in the center of the maze at the junction of the four arms and then allowing the mouse to freely explore the maze for 10 minutes.

### Fear conditioning

Fear conditioning tests were performed for 2 days (Panlab Harvard Apparatus, Barcelona, Spain). On the first day, the mouse was placed in the center of the chamber and subjected to a foot shock of 7 mA for 1 second for fear-conditioning. At 24 hours after fear learning, the mice were placed in the same chamber without a foot shocks for 5 minutes to measure contextual fear retrieval.

### Western blots

The sample extracted from hippocampus of control and material separation mice. The cells were homogenized in lysis buffer containing 25 mM Tris·HCl pH7.6, 150 mM NaCl, 1% NP-40, 1% sodium deoxycholate, 0.1% SDS, and protease inhibitors. The cell lysates (20 μg) were electrophoresis using SDS gels and transferred to nitrocellulose membranes and the incubated with TrkB antibody (1:1000, R&D systems, goat, AF1494), CRHR1 antibody (1:1000, GeneTex, goat, GRX88961), CRHR2 antibody (1:1000, ORIGENE, rabbit, AP17244PU-N), Grin2B antibody (1:1000, proteintech, rabbit, 21920-1-AP), HTR1B antibody (1:1000, Aviva system biology, rabbit, OAAF02797), DCX antibody (1:1000, Cell Signaling Technology, rabbit, #4604), MAP2 antibody (1:1000, Cell Signaling Technology, rabbit, #4542), SOX2 antibody (1:1000, Cell Signaling Technology, rabbit, #23064), NeuN antibody (1:1000, Cell Signaling Technology, rabbit, #24307), Erk antibody (1:1000, Cell Signaling Technology, rabbit, #4695), p-Erk(thy202/204) antibody (1:1000, Cell Signaling Technology, rabbit, #9101), mTOR antibody (1:1000, Cell Signaling Technology, rabbit, #2983), p-mTOR antibody (1:1000, Cell Signaling Technology, rabbit, #5536) and Δ-actin antibody (1:10000, Santa Cruz biotechnology, mouse, Sc-47778 HRP) for 16h at 4℃. After washing with 1X TBST buffer, the blots were incubated with anti-goat or anti-rabbit IgG (Thermo Fisher Scientific, USA). Band images were obtained by using a Molecular Imager ChemiDoc XRS+ (Bio-Rad, Hercules, CA, USA), and band intensity was analyzed using Image Lab TM software version 2.0.1 (Bio-Rad, Hercules, CA, USA).

### Single nucleus RNA seq

Briefly, frozen hippocampus of 3 mice in each sample group were dissociated using the Adult Brain Dissociation kit (Miltenyi Biotec, Germany), and after dead cell removal, 10x Chromium Single Cell 3’ Protocol (CA, US) was used to prepare a cDNA library with a target of 10,000 cells per sample. performed. And the prepared library was analyzed using an Illumina HiSeq X ten sequencing system (CA, US).

For bioinformatic analysis, raw base call files from the sequencer were generated into FASTQ files using ‘mkfastq’, and alignment, filtering, barcode counting, and UMI counting were performed with ‘cellranger’. Seurat 4.0.5 R package was used for the following analyses: data pre-processing, quality check, dimensionality reduction, graph-based cell clustering, the top expressed biomarker finding, gene-set enrichment (using gProfiler), finding differentially expressed genes among groups, identifying subtypes of cell composition, and cell population analysis of single nucleus RNA sequencing data.

Brain dissociation, single nucleus RNA sequencing and bioinformatic analyses were performed by Macrogen (South Korea).

### Whole genome sequencing (WGS)

To investigate somatic mutations, we compared the genomes of the hippocampus (100x depth coverage) and the genome of the mouse heart (30x depth coverage).

Genomic DNA extracted from the heart and the hippocampus was used to construct a next-generation sequencing (NGS) library using a TruSeq DNA polymerase chain reaction-free kit. Whole-genome sequencing was performed using the Illumina NovaSeq 6000 instrument. Thereafter, the bioinfomatic analysis continued through the following process: aligning using the GRCm39 reference (bwa-mem(38)), removing redundant reads based on align and coordinate (SortSam(39), MarkDuplicate(39)), analyzing variations (single nucleotide polymorphism, insertion and deletion (bcftools)(40), structure variant (Delly(41)), copy number variation (CVNcaller(42)), and somatic mutation (VarScan2(43))), and annotating each variation (SnpEff, ANNOVAR(44)).

### Demographic data

This study was approved by Institutional Review Board of Inha University Hospital (2022-09-044); it was conducted between April and June 2022.

A web-based self-reported assessment was conducted for the community population self-quarantined following contact with patients with COVID-19, and a web-based survey was conducted upon obtaining consent. The information obtained from the study participants included sex, age, marital status, education, residence type, degree of depression (patient health questionnaire-9, PHQ-9), degree of anxiety (general anxiety disorder7, GAD7), somatic symptoms (patient health questionnaire-15, PHQ15), suicidal ideation (P4 screener), stress perception (Perceived Stress Scale, PSS), post-traumatic stress symptoms (Korean version of the Impact of Event Scale, IES-K), sleep disorder symptoms (Korean version of the insomnia severity index, ISI-K), and self-esteem status (Rosenberg Self-Esteem Scale, RSES).

### Quantification and statistical analysis

Statistical data analysis was performed using R (version 3.5.1) and Prism 8 (GraphPad). Data normality was determined using a Shapiro-Wilk normality test. Normally distributed data were analyzed using Student’s t-test and analysis of variance (ANOVA), followed by post-hoc tests. Data that failed the normality test were analyzed using the Mann-Whitney U test. The ROUT method was used to exclude outliers with a Q factor of 5%, considering the differences among wild-type mouse populations, such as resilience.

## Acknowledgements

This work was supported by the intramural study from National Center for Mental Health in South Korea (R2022-C to Y.K.), a National Research Foundation of Korea (NRF) grant funded by the Korean government (MSIT) (NRF-2021R1C1C1003266 to Y.K.), and the Korea Health Technology R&D Project through the Korea Health Industry Development Institute (KHIDI), funded by the Ministry of Health & Welfare, Republic of Korea (HI22C0492 to Y.K.). The authors declare that they have no competing financial interests.

## Author contributions

B.L. performed mice behavioral tests, immunoblotting, and genomic sequencing analysis. S.J. performed mice behavioral tests. S.I. performed genomic sequencing analysis. Y.S. and J.N.B. performed the demographic study. B.L., H.J.C., J.N.B., and Y.K. designed the experiments and wrote the manuscript.

## Declaration of interests

There is no conflict of interest.

## Notes

### Competing Interest Statement

The authors have declared no competing interest.

