## Supplementary Figure 1 and 2 for "Translational approach to social isolation during a global pandemic: Hippocampal somatic mutation"

Genomic DNA extracted from the heart and the hippocampus was used to construct a next-generation sequencing (NGS) library using a TruSeq DNA polymerase chain reaction-free kit. Whole-genome sequencing was performed using the Illumina NovaSeq 6000 instrument. Thereafter, the bioinfomatic analysis continued through the following process: aligning using the GRCm39 reference (bwa-mem[18]), removing redundant reads based on align and coordinate (SortSam[19], MarkDuplicate[19]), analyzing variations (single nucleotide polymorphism, insertion and deletion (bcftools)[20], structure variant (Delly[21]), copy number variation (CVNcaller[22]), and somatic mutation (VarScan2[23])), and annotating each variation (SnpEff, ANNOVAR[24]).

### Supplementary Figures and Legends

**Supplementary Figure 1. Exploring corticotropin-releasing hormone receptor 1 (CRHR1)-related downstream signaling using immunoblotting**

(**A**) Compared to the control group (CTL), the maternal separation with social isolation group (MS+SS) showed significant increases in Microtube-associated protein 2 (MAP2). No difference was observed in the major downstream signaling and associated proteins of CRHR1, such as GRIN2B, mTOR, p-mTOR, ERK, and p-ERK. (n = 3 mice each group; 6 mice each group in MAP2 assay only. One-way analysis of variance, ** $p<0.01$, * $p<0.05$, post-hoc Tukey t-test)

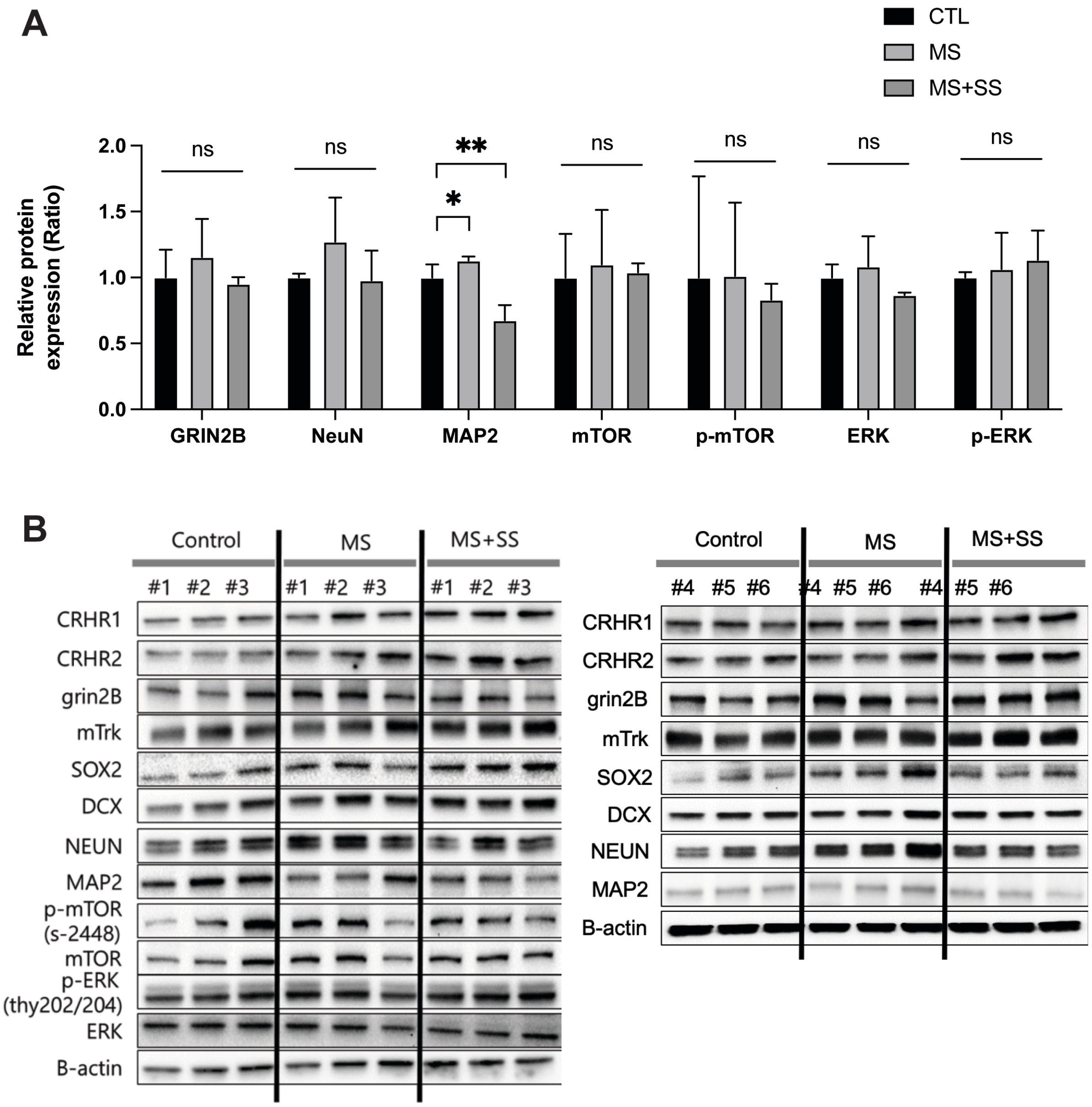
A
CTL
MS
MS+SS
Relative protein expression (Ratio)
2.0
1.5
1.0
0.5
0.0
ns
ns
**
*
ns
ns
ns
ns
GRIN2B
NeuN
MAP2
mTOR
p-mTOR
ERK
p-ERK
B
Control
MS
MS+SS
#1 #2 #3 #1 #2 #3 #1 #2 #3
CRHR1
CRHR2
grin2B
mTrk
SOX2
DCX
NEUN
MAP2
p-mTOR (s-2448)
mTOR
p-ERK (thy202/204)
ERK
B-actin
Control
MS
MS+SS
#4 #5 #6 #4 #5 #6 #4 #5 #6
CRHR1
CRHR2
grin2B
mTrk
SOX2
DCX
NEUN
MAP2
B-actin

**Supplementary Figure 2. Single nucleus transcriptomic characteristics including cell type distribution and gene expression patterns in the hippocampi of mice with maternal separation and social isolation**

(**A–D**) Identified cell types by Seurat analysis from single nucleus RNA sequencing in mouse hippocampi and cell type distribution between the groups. Graph theory-based cell type clustering of **C** is presented. The identified cell types classified in the control group (CTL) included neurons, which appeared the most at 92.1% compared to the maternal separation group (MS) and the maternal separation with social isolation group (SS).

(**E**) Heatmap showing the gene expression patterns for the top 20 genes in each cluster.

(**F**) Dot plot showing the top 20 genes per cluster. The size of each dot represents the percentage of the gene per each cluster. Color represents the average expression levels. (n = The hippocampi of 3 mice each were included in the CTL, MS, and SS).

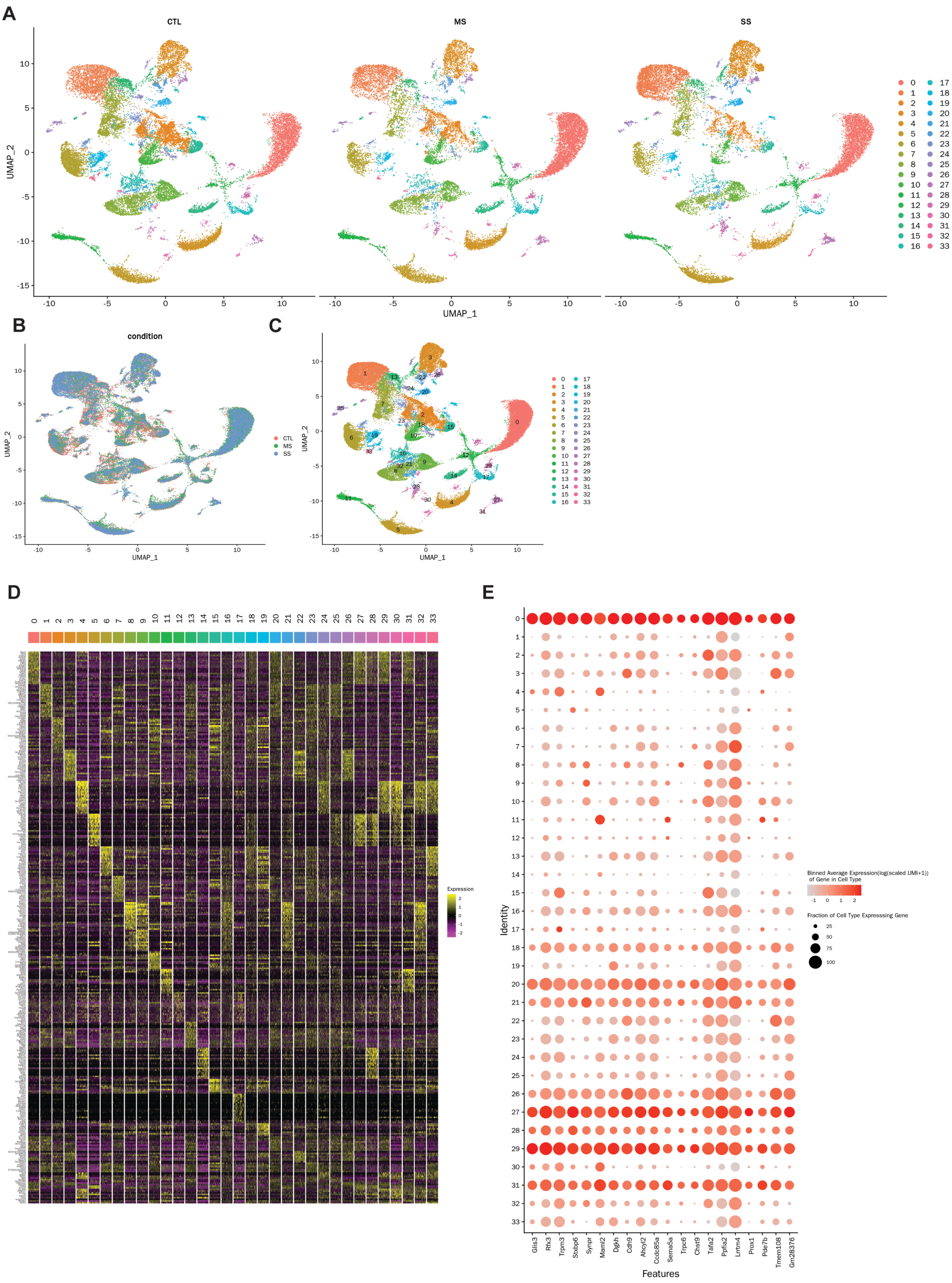

A
CTL
MS
SS
UMAP_2
UMAP_1
B
condition
CTL
MS
SS
C
D
Expression
E
Identity
Features
Glis3
Rfx3
Trpm3
Stxbp6
Synpr
Maml2
Dgkh
Cdh9
Ahcyl2
Ccdc85a
Sema5a
Trpc6
Chst9
Tafa2
Ppfia2
Lrrtm4
Prox1
Pde7b
Tmem108
Gm28376
Binned Average Expression(log(scaled UMI+1)) of Gene in Cell Type
Fraction of Cell Type Expressing Gene

**Supplementary Table 1. Somatic mutation profiles for each mouse individual.**
